# A novel metric reveals previously unrecognized distortion in the analysis of scRNA-seq data

**DOI:** 10.1101/689851

**Authors:** Timothy Hamilton, Breanne Sparta, Shamus Cooley, Samuel D. Aragones, J. Christian J. Ray, Eric J. Deeds

## Abstract

High-dimensional data are becoming increasingly common in nearly all areas of science. Developing approaches to analyze these data and understand their meaning is a pressing issue. This is particularly true for single-cell RNA-seq (scRNA-seq), a technique that simultaneously measures the expression of tens of thousands of genes in thousands to millions of single cells. Popular analysis pipelines significantly reduce the dimensionality of the dataset before performing downstream analysis. One problem with this approach is that dimensionality reduction can introduce substantial distortion into the data, particularly by disrupting the local neighborhoods of certain points. Since many scRNA-seq analyses like cell type clustering or trajectory inference rely on these near-neighbor relationships, distortion in this aspect of the data could significantly influence the outcomes of these analyses. Here, we introduce a straightforward approach to quantifying this distortion by comparing the local neighborhoods of points before and after dimensionality reduction. We found that popular techniques like t-SNE and UMAP introduce substantial distortion even for simple simulated data sets. For scRNA-seq data, we found the distortion in local neighborhoods was often greater than 95%, and that there was no consistent set of neighborhoods across the various steps in the consensus scRNA-seq analysis pipeline. We also found that this distortion had profound impacts on the outcomes of cell type clustering and other downstream analyses. Our findings suggest that caution must be applied when interpreting results in terms of 2-D visualizations produced by tools like UMAP, and that there is a critical need for new dimensionality reduction tools that more effectively preserve the local topological structure of the data.

## Introduction

The combination of computer science and molecular biology has empowered a revolutionary era for characterizing the biological processes that underpin how cells grow, mature and respond to their environment. For example, single-cell RNA-sequencing (scRNA-seq) data is notoriously high dimensional, consisting of counts of mRNA molecules for around 20,000 genes in tens of thousands to millions of individual cells. Here, classic approaches to manually or visually interpret patterns of gene expression across 20,000 genes is quite obviously out of the question^1–3^. As such, a robust ecosystem of computational tools has emerged to support the accessibility of single-cell technologies, including popular pipelines like Seurat and Scanpy^4,5^. On the surface, these computational tools may appear to resolve the challenges associated with understanding data of this complexity, but these tools also entail a large number of algorithmic choices that must be made to productively analyze the data. For example, the extremely high dimensionality of the data can make certain computations inefficient, thus taking unreasonable amounts of time to complete. Further, the large number of features in the data reduces the power of any statistical test due to multiple hypothesis testing. To overcome these challenges, which are often collectively referred to as the “curse of dimensionality,” the use of dimensionality reduction has become ubiquitous across various steps of single-cell analysis pipelines^2,6–8^.

The most common first step of this analysis pipeline involves “feature selection,” which identifies a subset of genes that are thought to be the most biologically informative for identifying cell types or characterizing cellular responses to perturbations^2–5^.This is typically done using the “Highly Variable Genes” (HVG) approach, which identifies genes in the data whose variance is higher than would be expected given a gene’s average expression level. While there are some rough guidelines available in the literature, the choice of the number of HVGs to use for downstream analysis is ultimately arbitrary; it is usually set to be around 2,000 or so^2,9^. This obviously represents a significant reduction in the dimensionality of the data. The data is then subjected to several nonlinear transformations, such as “Counts Per Million” or CPM normalization and log and z-score transformations^2,3,9^. After normalization, dimensionality reduction is typically performed using PCA, a linear approach, although non-linear approaches like autoencoders can be used for this step as well^3,10–13^. The choice of the number of dimensions to use here is again ultimately arbitrary, though one can use approaches like the “skree plot,” which attempts to find a breakpoint where explained variance does not significantly increase upon addition of further dimensions, or other approaches like molecular cross-validation^2,3,5,14^. Thus, while some guidelines for making these algorithmic cuts do exist, it is currently not well understood how relying on lower-dimensional representations of the data influences the outcomes of these studies.

Most downstream analysis of scRNA-seq data, including cell-type clustering, pseudotime or RNA velocity analysis, etc., is performed on feature-selected, CPM normalized, log-transformed, PCA-reduced data. After this analysis is performed, the data is often visualized in two dimensions using nonlinear dimensionality reduction tools like t-Stochastic Neighbor Embedding (t-SNE) or Uniform Manifold Approximation and Projection (UMAP)^2,3,15,16^. Much of the evaluation of these data, including tracking changes in gene expression for critical genes in disease states vs. healthy controls, or reasoning about changes in cell state during differentiation and development, is done using these 2-D visualizations^2,5,17,18^. Recent work, including our own, that has shown that these visualizations fail to accurately depict the high dimensional data they are meant to represent^17,19,20^.

In order to understand the issues that might be introduced through dimensionality reduction, consider the familiar problem of making a 2-D map of the entire surface of the 3-D Earth. Doing this requires “slicing” the earth along some axis to unfold it into a map; this is commonly done in a line through the Pacific, since few landmasses are disrupted by this cut. Then, the mapmaker must either increase the relative size of landmasses near the poles or slice the map again to project the globe into two dimensions. Regardless of technique, the globe cannot be represented in two dimensions without slicing and distorting the map in some way, which has led, for instance, to popular criticisms of the Mercator Projection. While distortion of distance and area are of course important, perhaps more concerning is the fact that the discontinuous slices mentioned above take points that are nearby (e.g., Alaska and Russia) and place them on opposite sides of the map. This means that the local neighborhoods of many of the points on the globe are completely different between the Earth itself and the 2-D representation.

The scale of dimensionality reduction in scRNA-seq is much larger, since analysis of that data is not merely 3-D to 2-D but rather 20,000-D to around 50 or even 2 dimensions. It has been reported that two-dimensional t-SNE and UMAP projections are significantly distorted, as the embedded representation does not coherently map to the structure of the original, high-dimensional data^17,19,20^. Yet whether the severe distortion arises during the production of a 2-D t-SNE or UMAP, or during the application of earlier steps in the single cell analysis pipeline, is not well understood. Also, it is not clear whether a consistent set of cell neighbors that map to functionally similar groups of cells can be reproducibly uncovered and preserved throughout the analysis pipeline. Answering these question entails systematically considering how each of the dimensionality reduction steps in the standard pipeline influences the structure of the data.

We approached this task by applying a simple statistic, inspired by the above metaphor of the globe, to quantify the extent to which any given dimensionality reduction technique discontinuously slices or folds the data in some way. This statistic is based on comparing the *local neighborhood* of a point in a dataset with the local neighborhood of that same point in the reduced-dimensional space using the Jaccard distance^20^ (Fig. 1C). To get a sense for this distortion across all the points in the dataset, we average this Jaccard Distance, resulting in the Average Jaccard Distance or AJD. We first applied our approach to the simple problem of embedding points on the surface of a hypersphere (which is a straightforward generalization of the sphere to more than three dimensions) into the appropriate latent dimension from a higher-dimensional space. When we quantified how popular algorithms represent data drawn from a known manifold structure, we found that many techniques, such as t-SNE and UMAP, not only introduced discontinuous slices into the data when trying to embed hyperspheres into two dimensions, but also when trying to embed into the correct latent dimension of the manifold. Indeed, we failed to identify any non-linear dimensionality reduction tool currently implemented in the scikit-learn package in Python that could successfully preserve the neighborhood relations of points from hyperspheres above approximately 10 dimensions. We found similar results for the output of popular tools for generating simulated scRNA-seq data sets; none of the popular nonlinear dimensionality reduction techniques could successfully embed any of the data, even in the known latent dimension.

**Figure 1.**
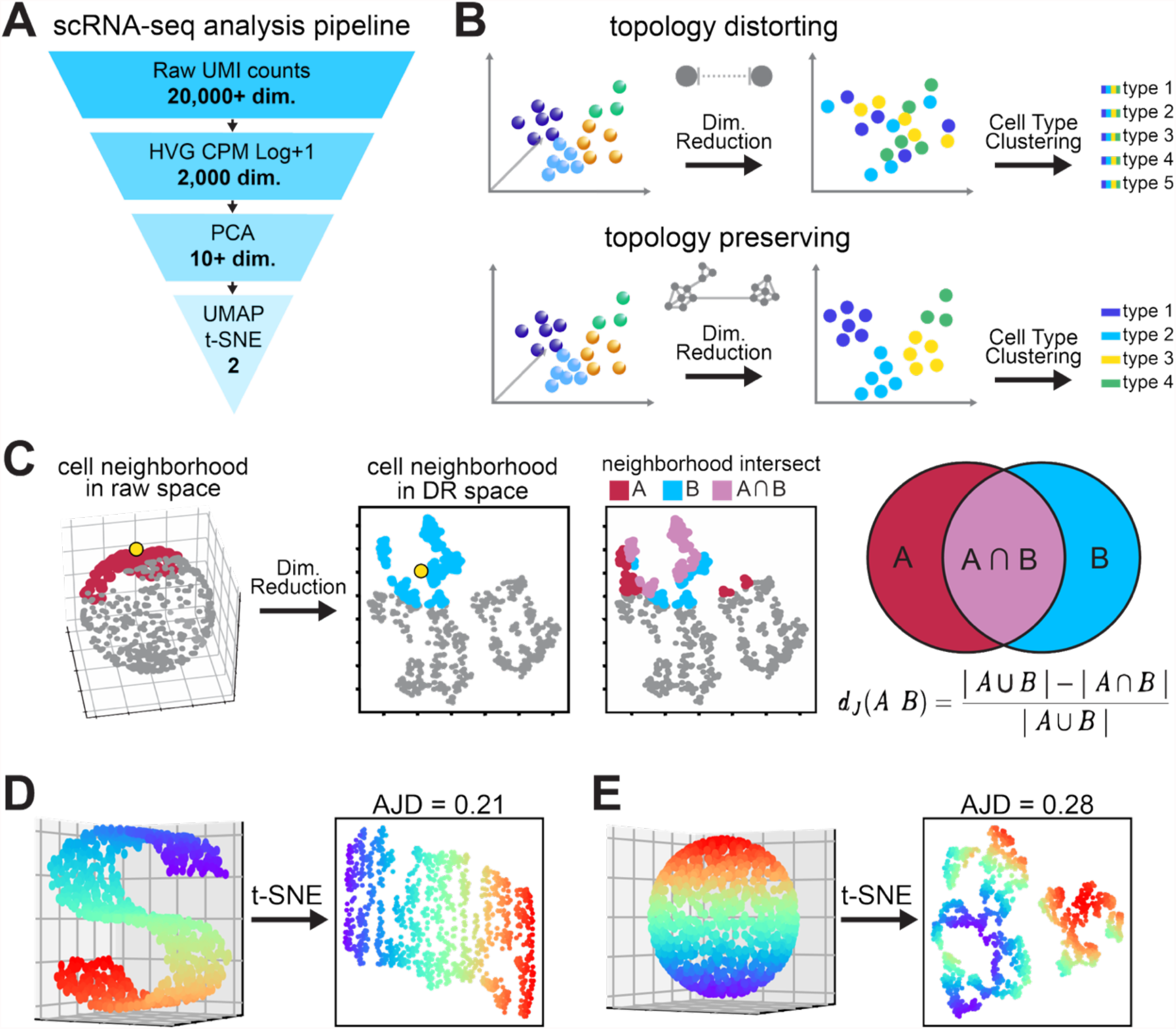
Dimensionality reduction and distortion. **A**. Flowchart of dimensionality reduction in the scRNA-seq pipeline, from ∼20,000 genes to ∼10-50 principal components, where clustering cells is generally done, to 2 UMAP/tSNE dimensions, where clusters are visualized, validated, and annotated. **B**. Schematic of topological distortion. If a transformation distorts topology, cell-type groups could be disrupted, resulting in problems with analysis or visualization. **C**. Schematic describing how we measured topological distortion. For a single point (shown in yellow) we find its *k*-Nearest Neighbors (kNNs), shown in red. Then we find the kNNs of that same point in the lower dimensional representation. We then compute the Jaccard Distance between them and average that distance across all the cells in the dataset to generate the AJD. **D**. Application of t-SNE to the S-curve dataset can unravel the 3D structure into a 2D plane. Note that this still introduces distortion, resulting in an AJD of 0.21 for *k* = 20. **E**. Application of t-SNE to a sphere, which cannot be globally embedded in 2-D. Note the larger number of discontinuous “tears” in the dataset compared to the S curve, which disrupts the rainbow pattern and leads to a larger AJD value of 0.28 for *k* = 20.

We then used our statistic to analyze how dimensionality reduction affects analysis of published scRNA-seq data. We carried out the “standard” scRNA-seq analysis pipeline on a total of 6 publicly available data sets and then, in a pairwise fashion, compared the AJD at each step in the process. Using our AJD metric, we found that each of these steps introduces tremendous distortion into the data. Specifically, commonly used pipelines disrupt <u>90-99%</u> of the local neighborhoods in the data when creating the representation that will be used for further quantitative analysis. This high AJD was consistent across all the steps we compared, indicating that there is no conserved neighborhood structure between the original data, the feature selected data, the normalized data, the PCA-reduced representation of the data used for clustering, and the data representation used for visual inspection. Like our findings on simulated data, even when embedding into higher dimensions, dimensionality reduction techniques generally introduced substantial discontinuity into the neighborhood structures. Interestingly, we found that the distortion is so severe that not only are *local* neighborhoods disrupted, but the *global* neighborhood structures are also severely permuted at every step of the standard pipeline. While PCA can find embeddings with relatively low levels of distortion, we found that this can only be achieved at much higher dimensionalities than are generally used for analysis.

Overall, our findings demonstrate that, regardless of the current techniques used to reduce dimensionality, the local and global structure of high-dimensional data is shuffled during each compression step that prepares scRNA-seq data for quantitative analysis. These observations suggest that any interpretation of lower-dimensional representations of scRNA-seq is dependent on the topological cuts induced during the analysis procedure. Rather than recovering a stable manifold of biological variation, we find that the results of cell-type clustering or psuedotime ordering can vary substantially depending on which algorithmic choices are being made. Our findings also suggest a need for new dimensionality reduction techniques that can reliably produce true topological embeddings, or at least closer approximations to manifolds of biological variation than current procedures can generate. We expect that the AJD statistic and the general approach introduced here will be helpful in evaluating and developing more effective tools for manifold learning and the effective analysis of scRNA-seq or other high-dimensional data.

## Results

### Developing a statistic to measure topological distortion

As discussed above, the “standard” scRNA-seq analysis pipeline entails several nonlinear transformations and dimensionality reduction steps, and it is currently unclear how each of these steps affects the structure of the data (Fig. 1A)^2,3,5^. Answering this question, however, requires defining the specific structural aspects of the data in which we are interested. Intriguingly, most scRNA-seq analyses rely on nearest-neighbor relationships, where neighborhoods are defined using the standard Euclidean distance in whatever space is being considered for the analysis^2,3,21^. For instance, cell-type clustering in scRNA-seq is essentially universally carried out by first determining the “*k*-Nearest Neighbors” (kNNs) of each cell, and using that information to build a reciprocal kNN graph. Clique detection algorithms like Louvain and Leiden are then used to cluster the resulting graph structure and determine the cell-type clusters in the data^2,3,5,22,23^. Many other analyses, like RNA velocity and pseudotime ordering, similarly rely on these nearest-neighbor relations^2,3,24,25^. As such, it is natural to ask to what extent these relationships are preserved during the analysis. From a mathematical perspective, this is equivalent to asking to what extent these transformations preserve *local topological* information.

Several approaches to measuring distortion due to dimensionality reduction have been proposed, but these focus primarily on the preservation of pairwise distances, not on the preservation of topological information. As such, inspired by the example of maps of the Earth described above, we developed an alternative statistic that focuses on how dimensionality reduction disrupts the local neighborhoods present in the data. To further motivate this, consider two simple datasets—the standard “S curve” (Fig. 1B) and a sphere (Fig. 1C). Algorithms like tSNE do a fairly good job of finding a 2-D embedding for the S curve, as represented by the preservation of the rainbow pattern in Fig. 1B. Application of t-SNE to a sphere, however, results in clear disruption of the rainbow pattern (Fig. 1C). To quantify the amount of topological distortion, we first take any given point of the dataset and compute its kNNs in the original dataset. So, if we take *k* to be 100, for any given point we will find its closest 100 neighbors in the original dataset. We can do the same in the lower-dimensional representation, which gives us two different sets of neighbors: the neighbors in the high dimensional data and the neighbors in the low dimensional data (the red and blue sets, respectively, in Fig. 1D). A very natural metric for comparing such sets is the “Jaccard distance” *d*_*J*_. If we call the original neighbor set “*A*” and the reduced-dimensionality neighbor set “*B*,” then this is defined as:

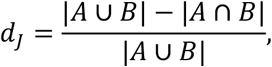

where |*X*| denotes the cardinality of the set *X*. The Jaccard distance is thus just the number of data points that are in the symmetric difference between *A* and *B*, which are the data points in *A* and *B* but not in both, divided by the total number of unique data points in either set. When this number is 0, it means the two sets are identical, and when it is 1, it means the two sets are disjoint. If the value is 0.5, this indicates that 50% of the combined set of neighbors are distinct between the lower-dimensional and higher-dimensional representation.

The above metric quantifies the distortion in the local neighborhood for any given point. To summarize distortion across an entire dataset, we generate a statistic that we call the “Average Jaccard Distance,” or AJD, which just averages the distortion for each point in the dataset. In other words, we calculate *d*_*J*_ independently for every point in the dataset, and then average those values to get the AJD. If we vary *k*, the AJD also provides a measure for how local the disruption of the data is.

### Application of AJD to Lower Dimensional Representations of synthetic data

The case of the sphere in Fig. 1C immediately suggests that nonlinear dimensionality reduction tools like UMAP and t-SNE can introduce significant distortion into lower-dimensional representations of data. We thus proceeded to systematically characterize this distortion for synthetic data sets whose properties are known *a priori*. A sphere in 3-D, for instance, cannot be represented in 2-D without introduction of discontinuities that break up local neighborhoods (as the example of 2-D maps of the Earth and Fig. 1D demonstrate). In this case, we chose to focus data sampled from a variety of structures: hyperspheres of varying dimensionality, multivariate Gaussian Distributions, as well as data from three synthetic scRNA-seq simulation approaches, Splatter, PROSST and ScDesign^26–28^. Our goal in each case was to generate a (relatively) high-dimensional dataset that contained a defined low-dimensional structure. To do this, we used a trivial approach to embed each dataset in a higher dimension by appending columns of 0s to the end of our lower dimensional data to reach the desired dimensionality. For instance, consider a hypersphere with latent dimensionality of 10, which would be referred to as *S*^9^ mathematically. A point in the dataset might look something like:

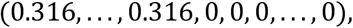

where first 10 coordinates are actual numbers specifying the position of a point on the surface of the hypersphere, and the remaining 90 coordinates are just 0. The fact that this manifold can be trivially embedded in 10 dimensions is clear, even by eye, but it is unclear if current dimensionality reduction tools can discover this fact.

As described in the Methods, it is relatively straightforward to sample from the surface of a hypersphere in any given dimension and then pad the dataset with the requisite number of 0s to generate a 100-dimensional vector. In Fig. 2A, we plot the AJD vs. embedding dimension for these test datasets using PCA, t-SNE and UMAP, three of the most popular dimensionality reduction tools currently in use in the community^3,11,12,15,16^. In each case, we compare the *k* = 20 nearest neighbors of each point in the original dataset and the embedding in the specified number of embedding dimensions, using the implementations of these tools in the scikit-learn package in Python^29^ applied to a hypersphere with 1000 points. PCA shows the expected behavior, with the AJD going to 0 at the minimum embedding dimension for each hypersphere. This is not surprising: these simple datasets only have *n* dimensions with any variation whatsoever (where *n* is the original embedding dimension of the hypersphere), and PCA of course identifies this. Surprisingly, however, the non-linear tools t-SNE and UMAP cannot solve even this simple problem. Indeed, for higher-dimensional datasets both t-SNE and UMAP fail to obtain an AJD less than 0.8, meaning 80% distortion of local neighborhoods, regardless of embedding dimension. Both tSNE and UMAP have free parameters, and for even the 20-dimensional hypersphere embedded in 100 dimensions, we could not find a combination of these parameters that resulted in low distortion for either algorithm (Supplementary Tables 2 and 3). Indeed, every single “manifold learning” algorithm implemented in the popular scikit-learn package in Python, including popular tools like IsoMap, Spectral Embedding and Multi-Dimensional Scaling (MDS), show similar results (Supplementary Fig. 1). Note that all of these results use the standard Euclidean distance to define the *k* Nearest Neighbors; using other metrics, like the *L*^1^ distance (i.e. the Manhattan distance), the *L*^∞^ distance (i.e. the Chebyshev distance), gives the same result (Supplementary Fig. 1). Also, increasing the number of points significantly (to 10,000, 50,000 or even 100,000) does not improve the performance of tolls like UMAP, and indeed, in some cases leads to even greater distortion (Supplementary Fig. 1).

**Figure 2.**
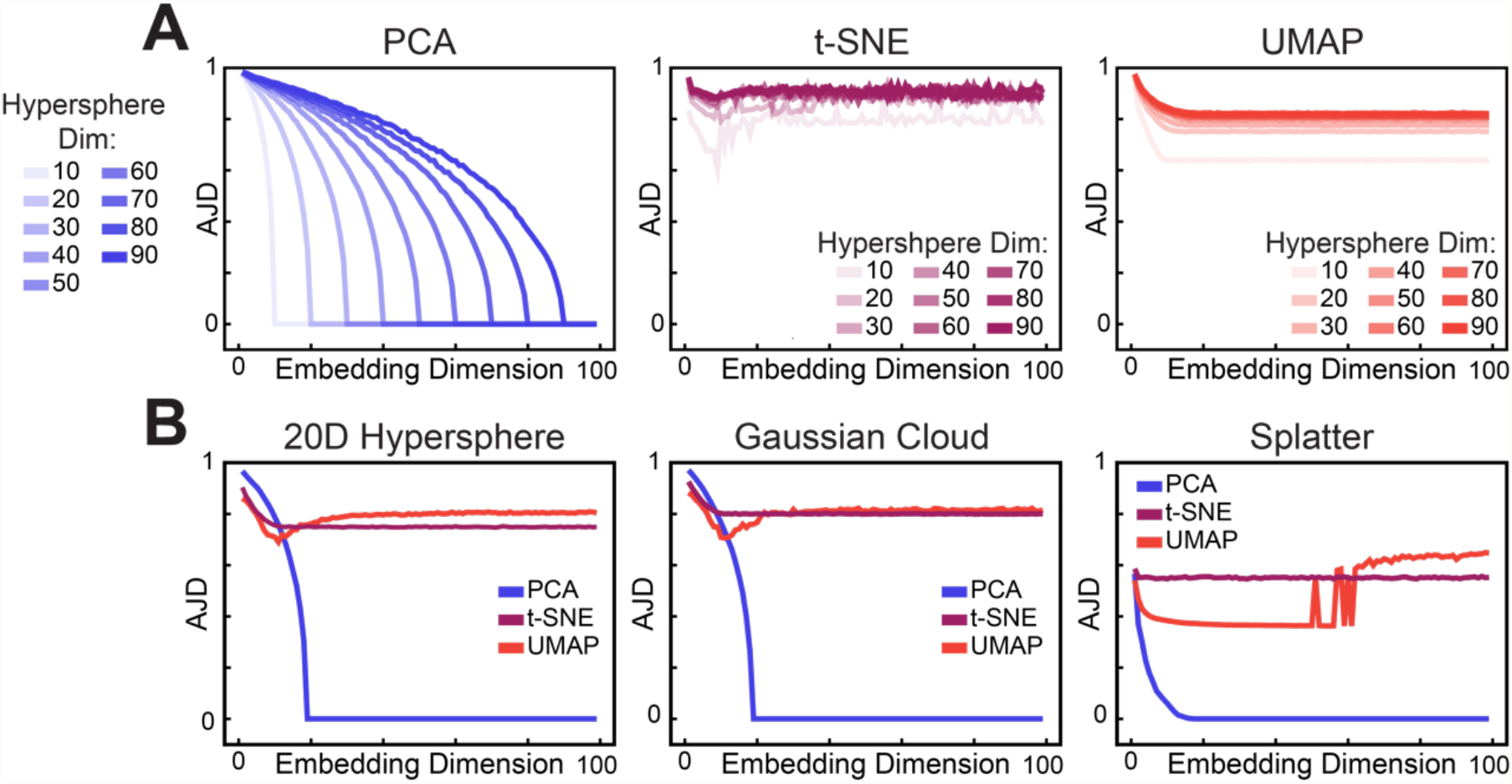
Applying dimensionality reduction techniques to synthetic data and measuring induced distortion with the AJD. **A**. AJD versus Embedding Dimension for PCA, t-SNE and UMAP applied to hyperspheres of given latent dimensions embedded in a 100-Dimensional space. Note for t-SNE, PCA was used to initialize the embedding,. **B**. AJD vs embedding dimension for PCA, tsNE and UMAP applied to various synthetic datasets. These are: a 20D Hypersphere embedded in a 100 Dimensional space, a 20D Multivariate Gaussian with identity-covariance embededed in a 100 Dimensional space and a mixture model made by the Splatter R-package, again embedded in a 100 Dimensional space.

In Fig. 2B, we compare PCA, t-SNE and UMAP for a hypersphere that can be embedded in 20 dimensions (this is technically the hypersphere *S*^19^). Again, neither t-SNE nor UMAP are able to obtain low distortion embeddings, even in 20 dimensions, while PCA can obviously find the appropriate embedding. We then applied this approach to additional datasets, starting with points sampled from a simple 20-dimensional multivariate Gaussian distribution. When we use the same approach to embed this structure in 100 dimensions, we again found that these nonlinear techniques cannot discover the trivial 20-D embedding of the data (Fig. 2B). We saw similar performance for simulated scRNA-seq data using Splatter (Fig. 2B), PROSST and ScDesign (Supplemental Figure 1B)^26–28^. Interestingly, across all of the tests shown in Fig. 2, t-SNE and UMAP generally show lower distortion than PCA in 2 dimensions, which is of course the dimensionality typically used for visualization. This suggests that these tools do actually preserve slightly more information when used for visualization, although it is important to note that the distortion is still very high.

### Evaluating Topological Distortion in scRNA-seq analysis

The above results suggest that dimensionality reduction techniques, and particularly t-SNE and UMAP, can indeed introduce high levels of distortion into the data. As mentioned in the Introduction, scRNA-seq analysis pipelines entail many different steps, all of which could in theory distort the data. We applied this pipeline to six different publicly-available scRNA-seq datasets (Supplementary Table 1). For each dataset, we compared the AJD between the following steps in the analysis: 1) the “Raw” data (i.e. the actual UMI count data obtained from the experiment itself); 2) after selecting 2000 “HVGs”; 3) “Transformed,” which entails CPM normalization with a scaling factor of 10,000, log-transformation (i.e. log (CPM + 1)), and z-score transformation; 4) “PCA,” which is performed on the transformed data to reduce dimensionality; 5) UMAP, which is applied to the PCA-transformed data and maps it to 2 dimensions. Note that we used the implementation of all of these tools in scanpy, a popular scRNA-seq analysis platform in Python, and we used visual inspection of the skree plot to choose the “elbow” dimension for PCA, as is generally done in the field^2,3,11,12^. Since we have five steps in this pipeline, we can calculate the AJD between each pair of steps. Here we chose *k* = 20 for our AJD calculations, resulting in the 5 × 5 AJD matrices shown in Fig. 3.

**Figure 3.**
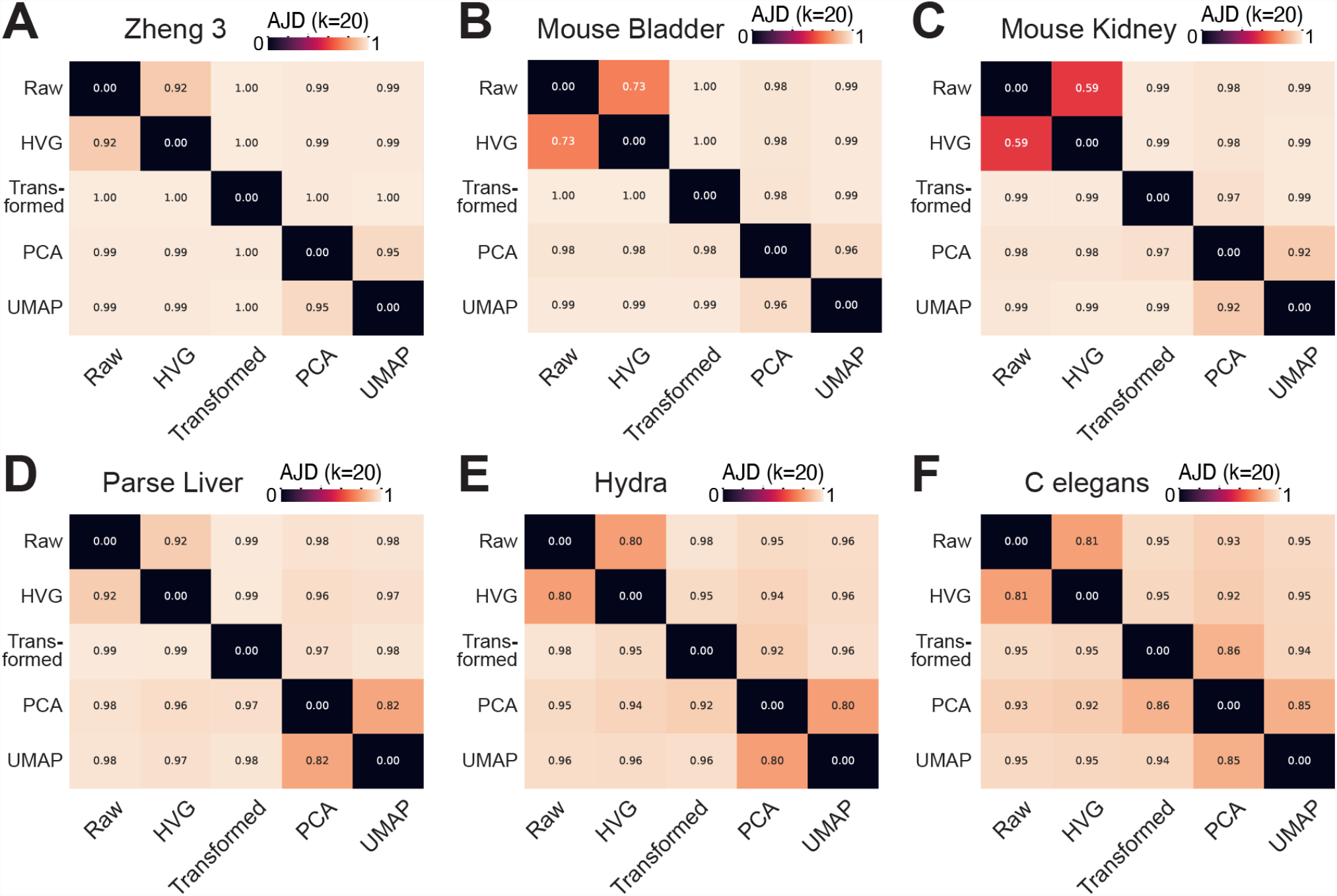
Pairwise comparison of the differences in neighborhood composition as measured by the AJD to determine the level of distortion introduced in various steps of the scRNA-seq pipeline. Listed are the following Datasets :**A**. Zheng 3 mixture of Monocytes, NK Cells and B-cells^30^ **B**. Mouse Bladder cells^31^ **C**. Mouse Kidney cells^32^ **D**.Parse Bio Liver cells^33^ **E**. Hydra cells^34^ **F**. Developing *C. Elegans* cells^35^

The first dataset we chose for this analysis is a classic human “Peripheral Blood MonoCyte” (PBMC) dataset consisting of a mixture of lymphocytes from Zheng *et al*^30^. The individual cells in this dataset were first separated into distinct cell-type groups (e.g. B cells, NK cells etc.) using FACS based on well-established cell surface markers, and as such this dataset has been used extensively in benchmarking studies for computational tools. We focused our analysis on what we call the “Zheng 3” dataset, which is the subset of B cells, NK cells and monocytes from Zheng *et al*. We also considered three datasets from mice, two of which were obtained using the 10X Genomics platform (Mouse Bladder and Mouse Kidney) and one of which was generated using the ParseBio platform (Parse Liver)^31–33^. We also included two datasets from non-mammalian sources: one whole-animal study based on the freshwater cnidarian Hydra and another from *C. elegans* embryos^34,35^. These datasets are summarized in Supplementary Table 1.

Interestingly, we see high levels of distortion no matter which two stages of the pipeline we are comparing (Fig. 3) across all datasets. The one exception to this is comparing the HVG data to the raw data: note here that no other transformations are applied, other than selecting the set of 2,000 HVGs. In some datasets, like the Mouse Bladder and Kidney, this results in only 59-73% distortion, which is the lowest we observed across all these calculations (Figs. 3B and C). In every other case, however, we observed AJD values very close to 1, suggesting that nearly 100% of neighborhoods are disrupted. Note that these extremely high levels of distortion are found regardless of which two steps in the pipeline are compared. This suggests that there is not a consistent set of neighborhoods that are preserved across any of the steps in the analysis. In other words, no two representations along the pipeline have the same *k*-nearest neighbor structure.

As mentioned above, in Fig. 3 we used the “elbow” dimension of the skree plot to determine the number of PCs for the “PCA” step of the pipeline, resulting in between 10 and 800 dimensions depending upon the dataset (see Supplemental Table 1). We next explored how changing the number of PCs changes the resulting AJD, comparing the embedding to the original Raw data (Fig. 4, purple curves) and the Transformed data (Fig. 4, green curves). Interestingly, the AJD between the Raw data and the PCA version was high regardless of embedding dimension, which is likely due to the fact that the AJD between the Transformed data, which is used as the input to PCA, and the Raw data itself is very high (Fig. 3). We can compare the PCA output to the Transformed data, and there we find that the AJD eventually reduces to close to 0, but only at dimensionalities well over 1,000, depending on the dataset (Fig. 4). This indicates that there is likely a lower-dimensional latent structure present in these transformed spaces, but the dimensionality of those structures is much, much higher than the number of dimensions typically used for analysis^2,3,5^. Interestingly, applying PCA to the Raw data directly identifies an even higher-dimensional structure, suggesting that the latent dimensionality of the structure one discovers depends critically on the transformations that are applied to the data (Supplementary Fig. 2).

**Figure 4.**
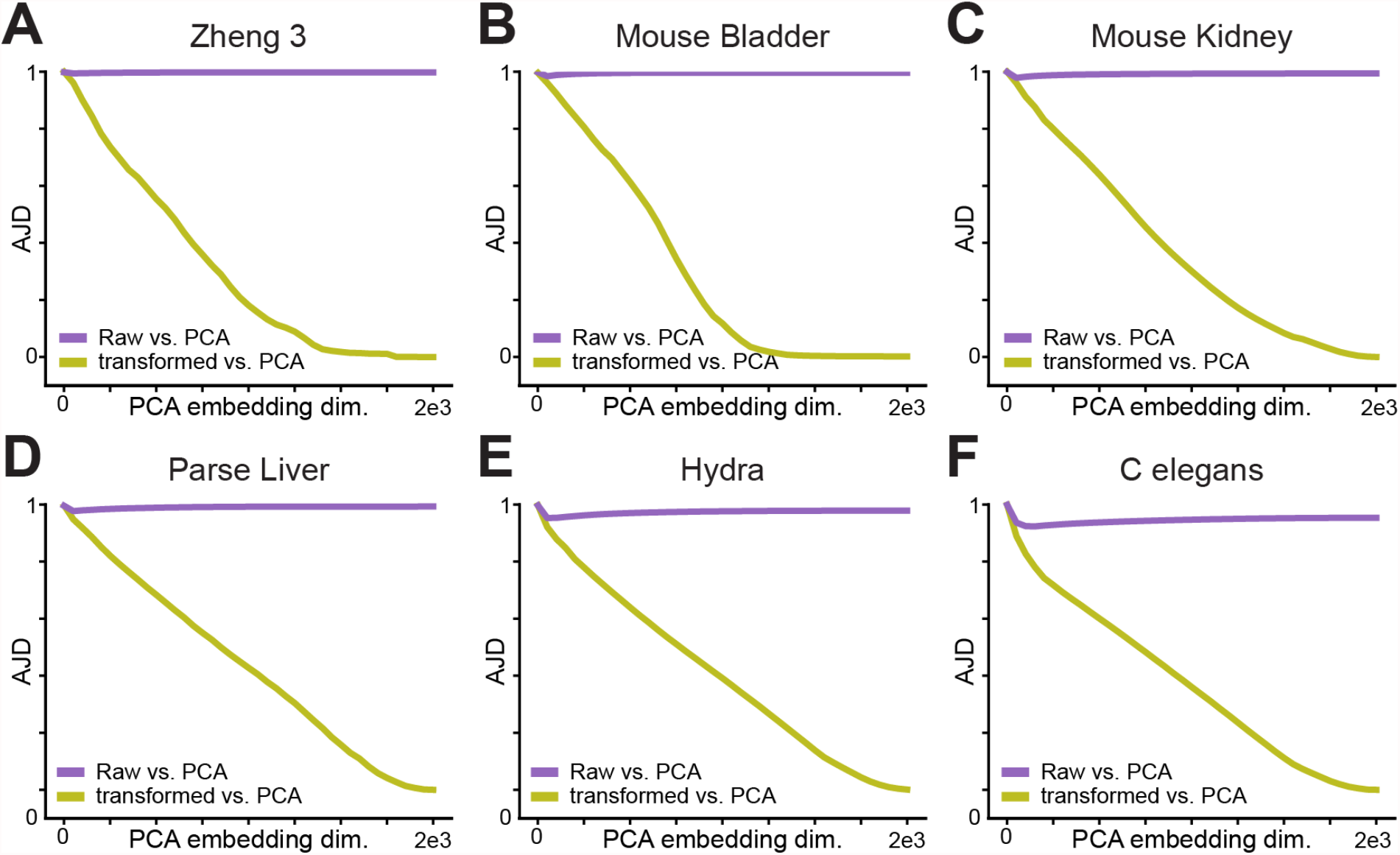
AJD vs embedding dimension for the main dimensionality reduction step of the scRNA-seq pipeline: Applying PCA to the “Transformed” data (Data after Feature Selection, CPM Normalization, and Scaling). The embeddings are compared to both the original Full transcriptomic “Raw” Space as well as the “Transformed Space. The specific datasets are: **A**. Zheng 3 mixture of Monocytes, NK Cells and B-cells^30^ **B**. Mouse Bladder cells^31^ **C**. Mouse Kidney cells^32^ **D**.Parse Bio Liver cells^33^ **E**. Hydra cells^33^4 **F**. Developing *C. Elegans* cells^35^

As mentioned above, non-linear dimensionality reduction algorithms have free parameters that can be chosen by the user; these include, for instance, the perplexity parameter for tSNE and the “n_neighbors” parameter for UMAP. We used the default values for those parameters for our results in Fig. 3, but one could also ask whether changing these parameters might improve the performance of the algorithm in terms of AJD^36^. To test this, we performed a simple grid search on the parameter space for tSNE and UMAP, focusing on the 2-D case since these algorithms are primarily used for visualization^2,3,17,36^. We applied this grid search to both the Zheng 3 and Hydra datasets, and found that, while optimization of the parameters could improve the AJD, the distortion was still quite high (Supplemental Tables 4 -7).

The results described so far have focused on a relatively small value of *k*, namely *k* = 20, for calculation of the AJD. While this is similar to the values typically used for scRNA-seq analysis (i.e. in clustering), the above results could represent relatively local permutations of the data. In other words, in the Zheng3 dataset, the distortion observed here could simply correspond to permuting the neighborhoods of B cells, but just to other B cells; such a local permutation might not introduce spurious relationships between, say, B cells and NK cells. To test this, we recomputed our comparisons for each of the scRNA-seq across various values of *k*. If the distortion is purely local, then AJD should decrease rapidly as *k* increases. What we found for the actual data was, however, quite the opposite: these “local” neighborhoods have to contain thousands of cells, often more than 50% of the dataset, for the AJD to decrease even below 0.5 (Fig. 5 and Supplementary Fig. 3). Note that this does not just hold for comparing the raw data to the final UMAP visualization; rather, distortion is highly non-local at each step in the pipeline.

**Figure 5.**
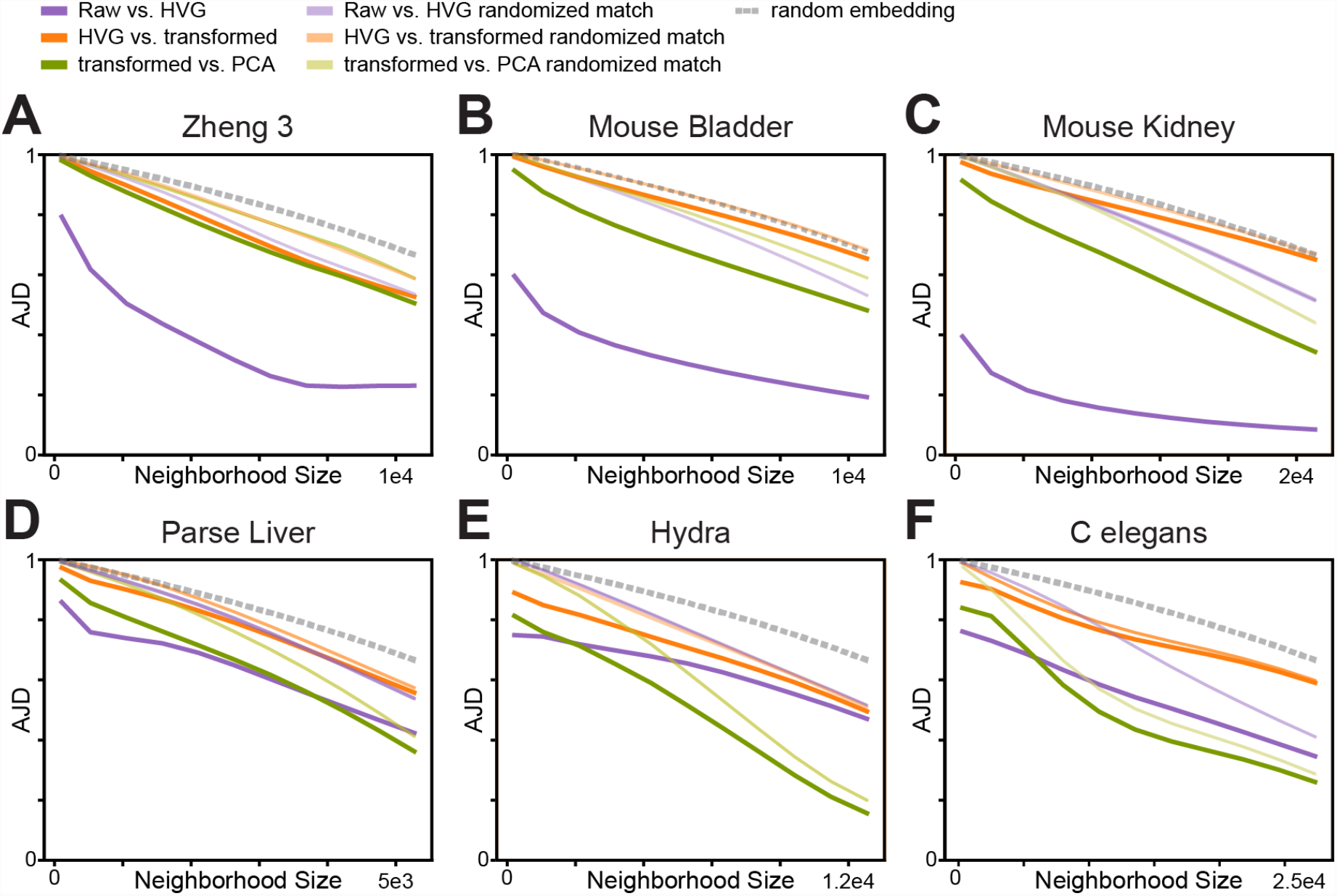
Comparison of how global the distortion injected into the data is, focusing on the comparisons of Raw vs HVGs, HVGs vs Transformed, and Transformed vs PCA. The comparison is carried out by computing the AJD repeatedly while varying *k*, the neighborhood size. For random controls, two separate experiments were done. First, in the more transparent lines, rather than the same cell’s neighborhoods in the the different spaces being compared, a random cell is chosen in the lower dimensional to compare its neighborhoods too. Second, in the dotted line, a graph meant to show the Expected Value of the AJD as a function of neighborhoods size if each cells neighborhoods in the lower dimensional representation was randomly composed of cells (see Supplementary Derivation 2 more detail). Listed are the following datasets: **A**. Zheng 3 mixture of Monocytes, NK Cells and B-cells^30^ **B**. Mouse Bladder cells^31^ **C**. Mouse Kidney cells^32^ **D**.Parse Bio Liver cells^33^ **E**. Hydra cells^34^ **F**. Developing *C. Elegans* cells^35^

The neighborhoods considered in Fig. 3 are so large that some degree of overlap could occur purely by random chance. We thus compared our results from the real data to a variety of random controls. The first such control is extremely simple, and considers the case where both versions of the dataset (i.e. the higher-dimensional and lower-dimensional versions) have neighborhoods constructed entirely at random. In other words, we assign each point a random set of *k* neighbors in the high dimensional space, and an independent random set of *k* neighbors in the lower dimensional state. We were able to calculate the AJD for this model analytically (see Supplementary Derivation 2), and while this simple “random embedding” model produced higher AJD vs. *k* curves than observed for the data, the results were surprisingly close (Fig. 3, dashed lines). As a second random control, we randomized the relationships between points in the high-dimensional data and the lower-dimensional representation. In other words, for any given comparison between two versions of the dataset, we first assigned each point in the high dimensional dataset to a random point in the low dimensional dataset. We then compared the neighborhoods of these high-dimensional points and their randomly chosen lower-dimensional counterparts to calculate the AJD for this case. Surprisingly, for most cases this result gives essentially the same AJD vs. *k* curve (Fig. 3).

Taken together, these results suggest that the distortion introduced by the various transformations and dimensionality reduction steps that are applied in the standard scRNA-seq analysis pipeline generate highly non-local distortion. In other words, these steps permute datasets in such a way that points that are in very different regions in the original data become close neighbors after the transformation. Moreover, these steps preserve about the same amount of topological information as one would expect if the points were assigned to one another randomly.

### Determining the effect of distortion on the results of scRNA-seq analysis

Our results above show that dimensionality reduction severely distorts the neighborhood structure of the data. The question then becomes: does this distortion impact the results of downstream analyses? For instance, if we perform clustering on the log-transformed HVG space and then on the PCA space, would we obtain the same sets of clusters? Since Louvain and Leiden, by far and away the two most popular clustering algorithms in scRNA-seq analysis, are both based on the nearest-neighbor graph, the tremendous change in the neighborhood structure described above could very well impact the clustering results^2,5,22,23^. To test this, we systematically compared the clustering results obtained from each step of the pipeline.

We applied this approach fist to the Hydra dataset, and found that the clusters produced by the Leiden clustering algorithm did indeed vary significantly between the Raw space, HVG-log-PCA space, and the UMAP space (Fig. 6 A-C). To quantify the extent of this disagreement, we used the Adjusted Rand Index, a metric that measures how similar two clusterings are relative to just randomly assigning cells to clusters. An ARI of 1 indicates perfect agreement between clusterings, while an ARI of 0 means the clustering results are essentially random. We noted that for both Louvain and Leiden that the number of clusters varied widely across the various spaces if we kept the resolution parameter the same (Fig. 6 A-C). As a result, we performed a simple grid search, calculating the ARI for every pair of resolution parameters and chose the number that gave the highest value (i.e. best agreement). In other words, say we compare clustering on the “transformed” and PCA versions of the Zheng 3 dataset. In each case, we tried a variety of resolution parameters and compared each set of clusters in the transformed space to each set of clusters in the PCA space and took the best overlap between these two spaces.

**Figure 6.**
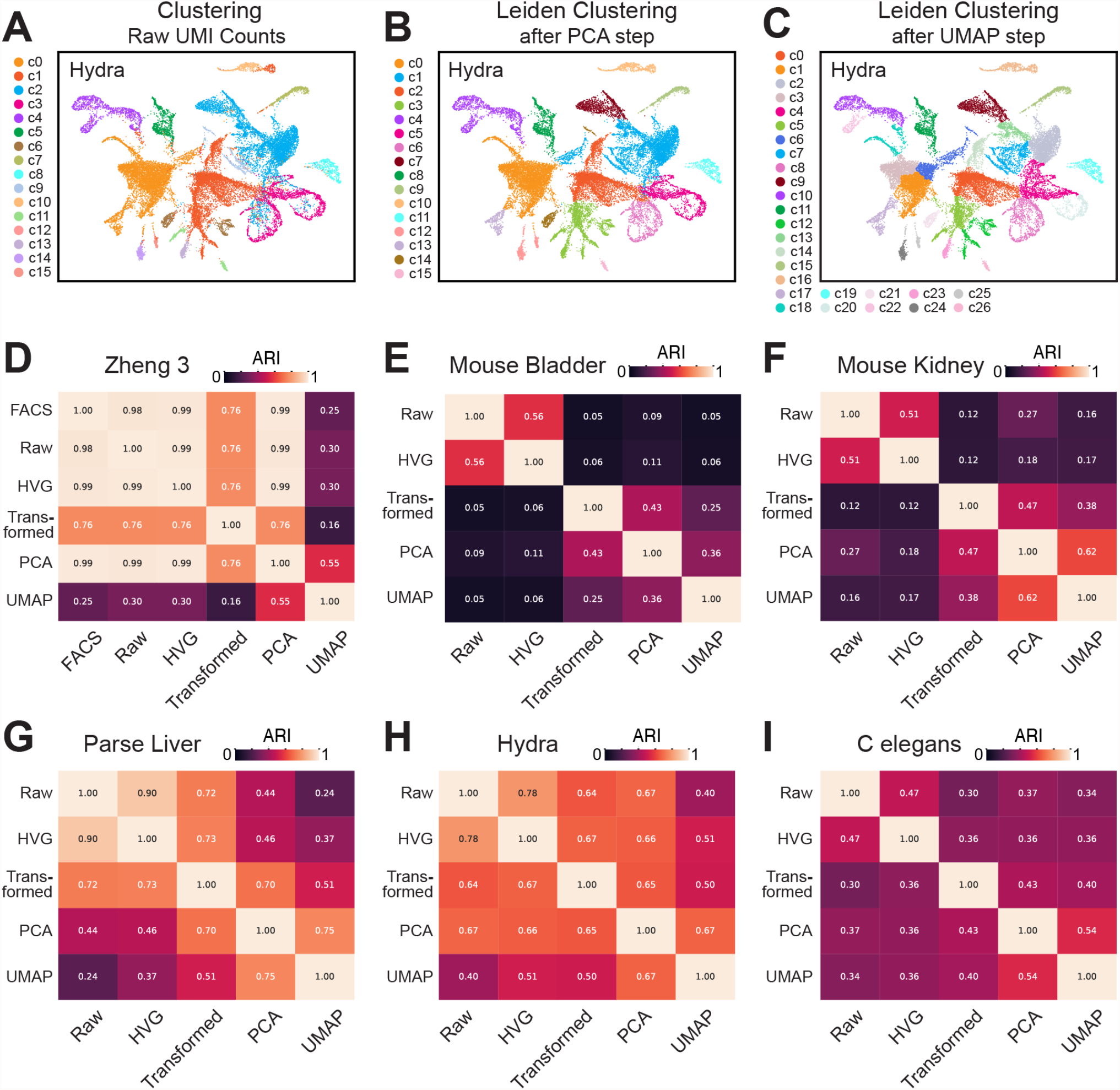
Downstream effects of distortion in scRNA-seq analysis. **A-C** Case study on how distortion effects clustering. Pictured in each panel is the Hydra scRNA-seq data with a computed UMAP projection. The spatial coordinates of each cell across each panel is the same; only the cluster assignment is different. The Panels are labeled by which space Leiden clustering is applied to using a resolution of 0.1. As shown, Clustering in differing spaces with the same resolution results in significantly different clustering results. **D-I**. Pairwise comparison of Leiden clustering for the six listed datasets analyzed. The ARI reported in each cell of the heatmap is the highest ARI reported between the Leiden clustering of the two different spaces as the resolution parameter was titrated.

The results of this calculation for the Leiden clustering algorithm are shown in the heatmaps in Fig. 6D-I. In the case of the Zheng 3 data, since we actually have the FACS labels, we can compare these clusters not only to one another, but also to the true biological identity of the cells. We see that clustering in the Raw, HVG and PCA spaces largely agrees with the FACS labels, while the transformed spaces, and *particularly* the UMAP space, does not agree nearly as well (Fig. 6D). Indeed, in the case of UMAP we see an ARI of 0.25 with the FACS labels, suggesting that this visualization does not represent these clusters very effectively. While the Zheng 3 data does show consistency across some of the steps in the pipeline in terms of clustering results, the other datasets we considered here do not. Indeed, we typically see ARI values between 0 (meaning completely random overlap) and 0.6 across transformations and datasets (Fig. 6 E-I). The results for the Louvain algorithm were essentially identical (Supplemental Fig. 4). Note that much of the agreement we observe requires careful tuning of the resolution parameter; just using the default value results in much worse overlap, even for the Zheng3 data (Supplemental Fig. 5). This suggests significant disagreement between the cluster structures obtained as the data is transformed and processed through the pipeline. This It is thus clear that, in addition to changing the structure of the data, there is not a consistent set of “cell identities” that are preserved through the pipeline. Rather, the massive structural changes that these steps entail generate rather different clustering results.

## Discussion

Dimensionality reduction is increasingly becoming a routine aspect of data analysis in scientific research. This is particularly the case in burgeoning field of single-cell transcriptomics, where much of the evaluation of the data rests on interpretation of these 2-D visualizations (e.g. coloring points in a UMAP based on expression of marker gene)^2,3,37,38^. While dimensionality reduction tools are highly popular, it remains unclear to what extent these approaches might introduce bias, artefacts or distortion as a part of computational analysis. Part of the issue in developing an understanding of these potential problems has been the fact that the vast majority of approaches to quantifying this distortion focus on how dimensionality reduction influences the pairwise distances between points^19,36^. While such distortion is interesting, in practice most analysis approaches tend to focus on near-neighbor relationships (e.g. clustering in scRNA-seq),where distortion will only have an effect if it impacts those local neighborhoods^2,3,24,39,40^.

Here, we applied a novel approach to quantifying distortion that focuses precisely on those local relationships. Specifically, the Jaccard distance is a metric that can directly compare the set of neighbors of a point in the high-dimensional space to the set of neighbors in the low-dimensional representation. Averaging this distance across all the points in the dataset allows us to characterize how much distortion in local neighborhoods is introduced by the dimensionality reduction process in question.

Surprisingly, application of the AJD statistic to a very simple test case revealed that none of the popular non-linear dimensionality reduction tools we tested can even solve the simple problem of finding the embedding of a hypersphere. This particularly concerning because the embedding of these hyperspheres in higher dimension was done completely trivially, by appending a string of 0’s to the end of the vector. Despite the fact that one can tell the correct embedding dimension of the hyperspheres purely by eye in this case, none of “manifold learning” tools we tested could discover the embedding. Thus, while these tools can perform fairly well (at least visually) on simple 2-D cases like the S-curve, they struggle with even simple manifolds in higher dimensions (Figs. 1 and 2).

Application of the AJD statistic to the scRNA-seq pipeline revealed that each of the steps in the pipeline introduce high levels of distortion into the data. In other words, the neighborhoods we find in the raw data are quite different from those present after HVG selection; log transformation, PCA and ultimately UMAP distort the data even further (Fig. 3). Further analysis showed that this distortion is not purely local; we see relatively high AJD values even when the “local neighborhoods” we consider contain thousands or tens of thousands of cells (Fig. 5). Interestingly, it is possible to introduce less distortion into the data when using PCA, but only when using many, many more principal components that are typically retained in an scRNA-seq analysis (thousands, compared to the tens to hundreds used in most analysis, Fig. 4).

One complication of this analysis is that is *a priori* unclear which of these levels of the steps in the pipeline should be taken as the “ground truth” for an AJD comparison. In other words, we might consider the nearest-neighbor relationships in the raw data to be the most meaningful: mathematical models that attempt to explain differentiation on the basis of Waddington’s landscape, for instance, work in the space of molecular concentrations, suggesting this might be the most meaningful space in which to search for cell types^41–45^. This is also the data directly generated by the experimental platform. Heuristic arguments have been made, however, to suggest that the standard pipeline de-noises and converts the data into a more meaningful state for analysis^2,3,30^. Regardless of these considerations, our results clearly show that there is no *consistent space* of neighborhoods across all these various representations; the raw, HVG, log-HVG, log-HVG-PCA and UMAP spaces are all quite different from one another. So, even if one takes the set of neighbors after the PCA space to be the “correct” or “most operationally useful” set of neighbors, the distortion introduced by the UMAP visualization is still between 80-90% (Fig. 3). In other words, the UMAP step introduces extremely high levels of distortion into the pictures that it generates, suggesting that the visual approaches used to validate applications of the pipeline might not be representative of the actual transcriptomic data itself.

Despite growing concerns regarding the over-application and over-interpretation of UMAP plots in single-cell genomics^17,19,46^, it is still quite common to read phrases to the effect of “analysis of the UMAP revealed…” in the recent literature (see, e.g., Li et al, Hozumi et al and Boutin et al)^44,47,48^. In other words, UMAP visualizations have clearly proven useful to researchers in this field, despite the fact that they preserve very little of the underlying structure of the data^47–51^. One potential explanation for this apparent contradiction is the fact that UMAP, and indeed the entire analysis pipeline, has a number of free parameters that can be varied in order to influence the outcome of the analysis. UMAPs can be re-generated indefinitely to achieve a visualization that captures some salient feature of the data; indeed, they can even be coaxed to produce arbitrary shapes like that of an elephant^17^. It could be that this flexibility allows researchers to find visualizations that, for instance, largely preserve the cluster structure they have generated. Such visualizations can then be further used to explore expression patterns that are (roughly) correlated with that cluster structure^47,49^. It could also be that the UMAP algorithm is somehow able to capture deeply salient features of the data and thus generated distorted images that are informative despite this distortion. UMAP itself was not explicitly designed to perform such information extraction; it was designed to find embeddings for manifolds, a task that our results on hyperspheres strongly suggest it struggles to complete effectively^15^. If UMAP is able to capture meaningful data from scRNA-seq, future work will be required to understand how it does this and what that means for the underlying structure of single-cell data.

Our work suggests that significant work needs to be done to develop new algorithms that can effectively embed high-dimensional manifolds. Recent that focuses on directly applying the mathematical definition of a manifold, namely as an atlas of local charts, have shown both theoretical and practical success^52^. Regardless, the AJD statistic discussed here holds clear promise for the development of approaches to both learn and visualize low-dimensional structures in data. In addition, our findings suggest that researchers should incorporate the AJD analysis into their pipeline, as this can allow them to evaluate the level of distortion they have in their own data, particularly to avoid overinterpretation of 2-D visualizations ^17,19,36^.

## Methods

### Average Jaccard Distance

To calculate the AJD, we find the *k* nearest neighbors for each data point *i* in the original ambient space and call this set *A*_*i*_. We similarly find the *k* nearest neighbors of that point *i* in the NDR-reduced space, which we call set *B*_*i*_. We used `sklearn.neighbors.NearestNeighbors`^29^ to calculate these nearest-neighbor sets, specifying the ball-tree algorithm due to its computational efficiency. To calculate the Jaccard distance between *A*_*i*_ and *B*_*i*_, we used the usual definition:

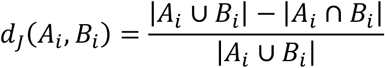

The AJD was calculated by taking the arithmetic mean of *d*_*j*_ across all points *i* in the dataset. While the analysis here is focused on measuring distortion between a higher- and lower-dimensional representation of a dataset, one could of course apply the AJD to any pair of representations of the same data for which the notion of *k* nearest neighbors is useful.

### Generation of Synthetic Datasets

#### Hyperspheres

The goal here was to sample *m* points uniformly from the surface of an *n* − 1-dimensional sphere in *d-*dimensional space. To do this, for each of the *m* data points we first sampled from a standard normal distribution *n* times (using the Python Numpy method `numpy.random.normal(0,1)`)^53^ to generate an *n*-dimensional vector. We then added enough 0s to the end of the vector to generate a *d*-dimensional vector and subsequently normalized the length of each sampled vector to 1, generating a uniformly-sampled dataset with the desired dimensionality.

#### Multi-dimensional Gaussians

The goal here was to sample *m* points from a *n*-dimensional multivariate gaussian in *d-*dimensional space. To do this, for each of the *m* data points we sampled from a standard normal distribution *n* times (using the Python Numpy method `numpy.random.normal(0,1)`). These samples became the first *n* coordinates of a vector. The remaining *n* + 1 to *d* coordinates were filled with zeros.

#### Splatter

To generate a mixture model via the `Splatter` package in R, we used the following steps^26^: First, we used the `splatSimulate` function, specifying (0.5, 0.25,0.25) as the probability of belonging in each of the 3 groups, 20 as the number of genes and 1000 as the number of points/cells. The other parameters used were set to the default for `splatSimulate`. To embed these simulated cells in 100-dimensional space, we added 80 zeros to the end of each simulated expression vector.

#### PROSST

To generate synthetic data meant to represent sampling from a temporal process (i.e cell differentiation), we used the PROSST package to generate a 20 dimensional dataset within a 100 dimensional space^27^. We first generated a random tree topology using the `gen_random_topology()` function with `n_branches` set to 9. The pseudotemporal distance between each branch point was set to 50. This tree structure was used to simulate a lineage with the `simulate_lineage()` function, with additional parameters “a” and “`intra_branch_tol`” set to 0.5, and “`nGenes`” set to 20. For this lineage, we then simulated the basal gene expression common throughout the pseudotemporal process using the `simulate_base_gene_expr`() function. We then computed two 20 dimensional vectors of random values. The first vector was sampled from a normal distribution with mean 0.2 and standard deviation 1.5 and the second vector was sampled from a normal distribution with mean 2 and standard deviation 1.5. These two vectors, along with the previously generated lineage were then used to generate a count matrix using the `sample_density()` function to sample 1000 cells, with resulting count matrix having a shape of 1000 cells by 20 synthetic genes. As above, a further 80 columns fully consisting of zeros were then padded to the end of the count matrix, resulting in a full dataset of 1000 cells and 100 features.

#### scDesign

To generate a mixture model via scDesign3 that was based upon sampling from real scRNA-seq data, we first loaded the single cell experiment data “`sce_filteredExpr10_Zhengmix4eq`” that was provided as part of the scDesign3 package, including the associated cell type information to form a single cell experiment object^28^. This single cell experiment object was then subsetted to only select the top 20 highly variable genes. This feature selected data, along with the cell type metadata, was then inputted into the `scDesign3` function, which uses a negative binomial model fitted to the top 20 genes with the annotated cell types provided as covariate information to generate a 1000 pseudocells, each with 20 pseudogenes. This data matrix was then padded with zeros to reach the final dimensions of 1000×100.

### Dimensionality Reduction

We executed dimensionality reduction with t-SNE, Isomap, Spectral Embedding, Multidimensional Scaling (MDS), and four versions of Local Linear Embedding (standard, modified, LTSA and Hessian Eigenmaps) using the standard implementations in the `scikit-learn` package in Python^29^. For UMAP we used `umap-learn`, which is a separate pacakge^15^. We implemented PCA using `sklearn.decomposition.PCA`, with the `svd_solver` set to “`arpack`”. For t-SNE, we used the implementation in `scikit-learn`, with in`it` set to “`pca`”, `random_state` set to 0 and `method` set to `exact`. For MDS, we used the implementation in `scikit-learn`, with `max_iter` set to 100 and `n_init` set to 1. For the nonlinear tools that had a `n_neighbors` parameter (UMAP, Spectral Embedding and Isomap), we set it to 100. For the linear tools that had a `n_neighbors` parameter, we set it to 101, except for Hessian Eigenmaps, for which we set n_neighbors to 231. For all other parameters not listed, we utilized the default values provided in `scikit-learn`.

### Literature scRNA-seq Datasets

To test the prevalence of distortion in real scRNA-seq data, we downloaded 6 cell × gene matrices from the publications listed in Supplemental Table 1. These matrices were generated by the original authors after the cell and gene filtering steps for doublets and empty droplets. These cell × gene matrices were saved as csv files to be later used in the single cell pipeline. We used all the cells available in these datasets, except in the case of the *C. elegans* dataset, in which only cells that were annotated with a definitive cell type were used. The resulting number of cells and genes are listed in Supplemental Table 1. These datasets are available on our GitHub site, https://github.com/DeedsLab/AJD_Metric.

### Standardized scRNA-seq pipeline

The six single cell datasets were then analyzed using `scanpy` following the tutorial posted to scanpy’s website. Each of the cell × gene matrices were read into a scanpy annodated data object via the pandas `read_csv` function and the Annotated Data object constructor^4^. This AnnData object was then used in the remainder of the analysis. We first normalized the data with the `normalize_total()` command, with the `target_sum` parameter set to 10000. Thus, while we refer to this as “CPM” normalization, in reality this is Counts per Ten Thousand normalization, as is current general practice in the field. We then log(*x* + 1) transform the data with the `log1p` command. Next, we selected the top 2000 variable genes using the `highly_variable_genes` function. These genes are then used to subset the original dataset, resulting in a purely feature selected dataset with no transformations termed “HVG”. The AnnData object was then scaled using the `scale()` command with the `max_value` parameter set to 10 as in the scanpy tutorial. This resulted in another dataset that we saved for analysis, termed “Transformed”. A PCA embedding was then calculated. First, all the possible components are calculated using the `pca()` command, with the solver set to “`arpack`” and the number of components set to 1999. Then, the kneedle algorithm implemented in the `kneed` package was used to find the number of pcs to use for the embedding, with each PC’s explained variance ratio passed into the `kl()` function of `kneed`, along with the parameter values of “`S`” being set to 1.0, curve set to “`convex`” and direction set to “`decreasing`”^54^. Once the number of PCs was determined the `pca()` command was called again, with the solver set to “`arpack`” and the number of principal component set as the number determined by the kneedle algorithm. The PCA embedding was then extracted and saved separately, which we termed “PCA”. Then, the nearest neighbors of the PCA embedding were found using the `neighbors()` function, with *k* set to 20. Then a UMAP visualization was constructed first by finding a PAGA embedding using the `paga()` function, with default parameter values. This PAGA embedding was used as the initial positions for the `umap()` function in scanpy, with the `init_pos` argument set to “`paga`”. This produced an embedding termed “UMAP” that was saved for further analysis.

### Randomized Control for AJD across pipeline steps

We created two different random controls to serve as reference for how local the topological distortion injected into the embedding by the scRNA-seq pipeline is. The first and more extreme control involves constructing one neighborhood for each cell, the simulated “high-dimensional” neighborhood, completely at random from all the cells in the dataset. We then created a similar, completely random “low-dimensional” neighborhood for each point. In other words, this control imagines that the neighbors of a point in the higher and lower dimensional versions of the data are assigned completely randomly. It is straightforward to derive the approximate expected AJD for this case analytically. The result is:

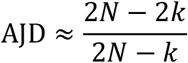

A detailed derivation of this equation is provided in the “Supplemental Derivation” PDF. This curve is plotted as the dashed line in Fig. 5.

While simple, the above equation is both approximate and does not respect any potential geometric constraints; in other words, it might not be possible to construct all the neighborhood relationships between real points using a meaningful metric in a finite-dimensional vector space. To deal with these issues, we generated a second kind of control, where we directly randomized the data in real examples of embeddings. To do this, we mapped each point in the high dimensional data to a randomly chosen point in the lower-dimensional representation. So, instead of comparing *A*_*i*_ and *B*_*i*_ to calculate the Jaccard distance for point *i*, here we compare *A*_*i*_ to *B*_*j*_, where *j* is some randomly chosen point from the lower-dimensional dataset. To compute these “random match” curves in Fig. 5, we took each successive step in the pipeline (i.e. Raw vs. HVG, HVG vs. transformed, etc.) and generated 100 independent random assignments of points in the higher-dimensional version of the data to points in the lower-dimensional data. We then averaged the AJD for these 100 random mappings at each value of *k*. This random control was done for each successive pair of steps in the pipeline down to the “PCA” step.

### Adjusted Rand Index

The Rand index quantifies the similarity between clusters in two partitions *U* and *V* (say, cell clusters in the ambient dimension and in a reduced dimension) through a contingency table that classifies pairs of points into four cases: pairs in the same cluster in both partitions (*a*), pairs in the same cluster in *U* but not *V* (*b*), pairs in the same cluster in *V* but not *U* (*c*), or pairs in different clusters in both partitions (*d*). It takes a value between 0 and 1. The Adjusted Rand Index (ARI) corrects the value by accounting for coincidental/chance clustering and avoiding the tendency of the unadjusted Rand index to approach 1 as the number of clusters increases. A detailed discussion of the ARI and it’s derivation may be found in ref. ^55^. We used the `adjusted_rand_score()`implemented in the `metrics` package in `scikit-learn` to compute the ARI values reported in this work.

### Clustering Comparison

To accurately measure how topological distortion affects the downstream analysis of each scRNA-seq dataset, we decided to measure how clustering results changed across each of the steps of the analysis, each of the steps were loaded into a separate AnnData object. Then for each of the separate steps of the pipeline, clustering was done in the following manner. First, a nearest neighbor graph was made in the appropriate space using scanpy’s `neighbors()` function, with the number of neighbors set to 15, and `n_pcs` set to 0 to force the function to use the appropriate space. Then, either Leiden or Louvain clustering was done using scanpy’s `leiden()` and `louvain()` command multiple time using resolution parameters ranging from 0.1 to 4.0 in steps of 0.5, resulting in 10 different clustering schema for each step of the pipeline. We then computed the ARI between each unique pair of clustering schema, and chose the pair with the maximum ARI value (and thus the greatest agreement between the two sets of clusters). These maximum ARI values are reported in Fig. 6.

## Supporting information

Supplemental Derivation

