## Supplemental Figures for "A novel metric reveals previously unrecognized distortion in the analysis of scRNA-seq data"

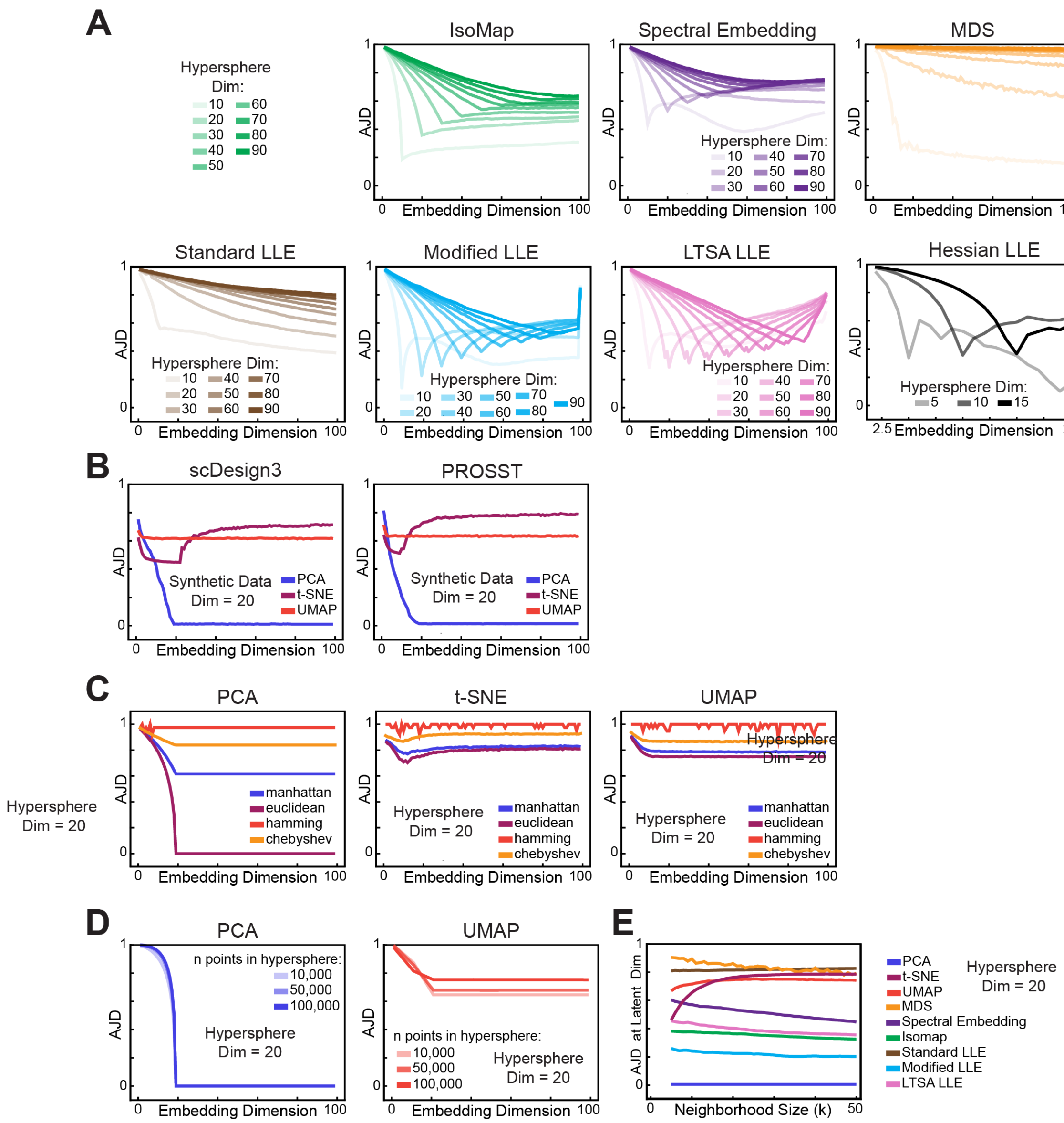


**Supplemental Figure 1.** Evaluation of distortion introduced by common dimensionality reduction techniques on synthetic data. **A.** AJD vs Embedding dimension for IsoMap Spectral Embedding and MDS on hyperspheres of various dimensions placed in an 100 dimensional space (20 dimensional space for Hessian LLE). **B.** AJD vs Embedding dimension for PCA, t-SNE, and UMAP as applied to different synthetic data meant to emulate the structure of scRNA-seq data. Shown here is the results from generated from scDesign3 and PROSST. In both cases, 20 genes were simulated and placed within a 100 dimensional space **C.** AJD vs Embedding dimension for PCA, t-SNE and UMAP applied to a 20 dimensional hyperspheres in a 100 dimensional space. The neighborhoods were calculated using the listed distance metrics (Manhattan (L1-norm), Euclidean (L2-norm), hamming, and Chebyshev distances). **D.** AJD vs Embedding dimension for PCA and UMAP applied to hypersphere from which the listed number of points were sampled from. **E.** AJD vs local neighborhood size for a 20 dimensional hypersphere in a 100 dimensional space that is embedded into a 20 dimensional space using PCA, t-SNE, UMAP, MDS, Spectral Embedding, Isomap, Standard LLE, Modified LLE, and LTSA LLE.

| Dataset Name | No. Cells | No Genes | Num PCs | GEO Accession No. |
|---|---|---|---|---|
| Zheng 3-mix | 21082 | 32740 | 12 | SRP073767 |
| Mouse Bladder | 20000 | 31055 | 26 | GSE163029 |
| Mouse Kidney | 43745 | 16275 | 39 | GSE107585 |
| Parse Liver | 9934 | 33059 | 22 | N/A |
| Hydra | 24459 | 17091 | 49 | GSE121617 |
| C.Elegans | 54649 | 20223 | 77 | GSE126954 |

**Supplemental Table 1.** Table showing information about each of the six real scRNA-seq datasets used in this study. This includes the name used in the paper, the number of cells and genes left after preprocessing and quality control, the number of principal components selected by the *kneed* package during the running of the pipeline and the GEO accession number if available.

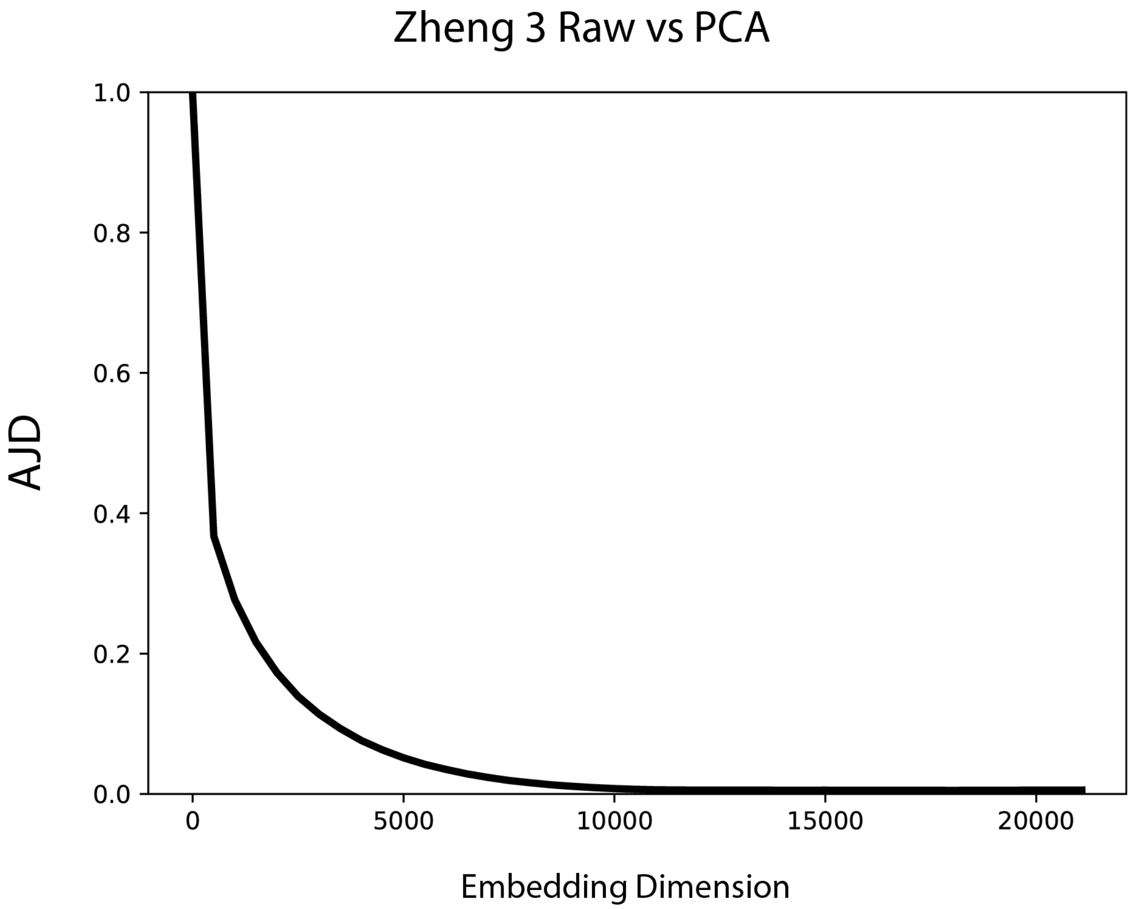


**Supplemental Figure 2.** Average Jaccard Distances versus Embedding dimension for the Zheng 3 dataset. Here, PCA was used to directly embed the original full data matrix to representations in varying dimensions, with 10,000+ dimensions required to ensure the embeddings have little topological distortion. This highlights that PCA can construct embeddings with little distortion, but only if a significant number of components are utilized, much larger than the number typically used in scRNA-seq data analysis.

| | **Minimum Distance** | | | | | |
|---|---|---|---|---|---|---|
| **k-Nearest Neighbors** | **0.0** | **0.1** | **0.25** | **0.5** | **0.8** | **0.99** |
| **10** | 0.831 | 0.815 | 0.787 | 0.747 | 0.705 | 0.680 |
| **20** | 0.800 | 0.781 | 0.746 | 0.689 | 0.631 | 0.597 |
| **50** | 0.785 | 0.760 | 0.730 | 0.680 | 0.610 | 0.582 |
| **100** | 0.778 | 0.754 | 0.720 | 0.667 | 0.602 | 0.569 |
| **200** | 0.768 | 0.743 | 0.711 | 0.651 | 0.587 | 0.552 |
| **400** | 0.759 | 0.731 | 0.702 | 0.638 | 0.573 | 0.534 |

**Supplemental Table 2.** Table showing the results of a grid search of the parameters for UMAP (k nearest neighbors and minimum distance). In each case we sampled 1000 points from a 20-dimensional hypersphere and embedded those points in 100 dimensions as described in the main text. The AJD shown is between the original 100 dimensional dataset and the version reduced down to 20 dimensions with UMAP using the listed parameters.

| | Perplexity | | | | | |
|---|---|---|---|---|---|---|
| **Learning Rate** | **5** | **25** | **50** | **100** | **200** | **400** |
| **12.5** | 0.827 | 0.791 | 0.749 | 0.697 | 0.694 | 0.708 |
| **25** | 0.825 | 0.719 | 0.691 | 0.630 | 0.634 | 0.566 |
| **50** | 0.845 | 0.779 | 0.747 | 0.695 | 0.632 | 0.628 |
| **100** | 0.868 | 0.805 | 0.780 | 0.726 | 0.645 | 0.566 |
| **200** | 0.882 | 0.914 | 0.809 | 0.741 | 0.670 | 0.647 |
| **400** | 0.948 | 0.906 | 0.866 | 0.735 | 0.642 | 0.530 |

**Supplemental Table 3.** Table showing the results of a grid search of the parameters for tSNE (learning rate and perplexity). In each case we sampled 1000 points from a 20-dimensional hypersphere and embedded those points in 100 dimensions as described in the main text. The AJD shown is between the original 100 dimensional dataset and the version reduced down to 20 dimensions with tSNE using the listed parameters.

| | **Minimum Distance** | | | | | |
|---|---|---|---|---|---|---|
| **k Nearest Neighbors** | **0.0** | **0.1** | **0.25** | **0.5** | **0.8** | **0.99** |
| **10** | 0.928 | 0.936 | 0.944 | 0.952 | 0.956 | 0.959 |
| **20** | 0.936 | 0.941 | 0.948 | 0.953 | 0.958 | 0.957 |
| **50** | 0.945 | 0.947 | 0.951 | 0.955 | 0.957 | 0.958 |
| **100** | 0.949 | 0.951 | 0.953 | 0.956 | 0.958 | 0.959 |
| **200** | 0.953 | 0.954 | 0.956 | 0.958 | 0.959 | 0.960 |
| **400** | 0.955 | 0.956 | 0.957 | 0.958 | 0.960 | 0.960 |

**Supplemental Table 4.** Table showing the results of a grid search of the parameters for UMAP (k nearest neighbors and minimum distance) that was applied to the Zheng 3-mix data at the PCA step. The AJD shown is between the PCA embedding of the transformed data and a two-dimensional UMAP embedding generated with the selected set of parameters.

| | Minimum Distance | | | | | |
|---|---|---|---|---|---|---|
| K-Nearest Neighbors | 0.0 | 0.1 | 0.25 | 0.5 | 0.8 | 0.99 |
| **10** | 0.812 | 0.801 | 0.799 | 0.807 | 0.837 | 0.844 |
| **20** | 0.814 | 0.806 | 0.804 | 0.812 | 0.831 | 0.840 |
| **50** | 0.833 | 0.822 | 0.820 | 0.815 | 0.837 | 0.841 |
| **100** | 0.844 | 0.833 | 0.823 | 0.827 | 0.838 | 0.848 |
| **200** | 0.853 | 0.839 | 0.827 | 0.831 | 0.843 | 0.851 |
| **400** | 0.858 | 0.849 | 0.838 | 0.837 | 0.848 | 0.860 |

**Supplemental Table 5.** Table showing the results of a grid search of the parameters for UMAP (k nearest neighbors and minimum distance) that was applied to the Hydra data at the PCA step. The AJD shown is between the PCA embedding of the transformed data and a two-dimensional UMAP embedding generated with the selected set of parameters.

| | **Perplexity** | | | | | |
|---|---|---|---|---|---|---|
| **Learning Rate** | **5** | **25** | **50** | **100** | **200** | **400** |
| **12.5** | 0.854 | 0.865 | 0.885 | 0.908 | 0.926 | 0.941 |
| **25** | 0.850 | 0.842 | 0.860 | 0.889 | 0.915 | 0.935 |
| **50** | 0.850 | 0.831 | 0.841 | 0.869 | 0.903 | 0.930 |
| **100** | 0.852 | 0.827 | 0.829 | 0.851 | 0.890 | 0.925 |
| **200** | 0.852 | 0.827 | 0.826 | 0.840 | 0.880 | 0.921 |
| **400** | 0.852 | 0.827 | 0.825 | 0.833 | 0.871 | 0.918 |

**Supplemental Table 6.** Table showing the results of a grid search of the parameters for tSNE (learning rate and perplexity) that was applied to the Zheng 3-mix data at the PCA step. The AJD shown is between the PCA embedding of the transformed data and a two-dimensional tSNE embedding generated with the selected set of parameters.

| | **Perplexity** | | | | | |
|---|---|---|---|---|---|---|
| **Learning Rate** | **5** | **25** | **50** | **100** | **200** | **400** |
| **12.5** | 0.751 | 0.751 | 0.763 | 0.777 | 0.797 | 0.810 |
| **25** | 0.746 | 0.734 | 0.746 | 0.763 | 0.781 | 0.795 |
| **50** | 0.745 | 0.724 | 0.732 | 0.750 | 0.766 | 0.781 |
| **100** | 0.747 | 0.718 | 0.722 | 0.737 | 0.755 | 0.771 |
| **200** | 0.749 | 0.715 | 0.715 | 0.727 | 0.746 | 0.763 |
| **400** | 0.749 | 0.714 | 0.711 | 0.720 | 0.739 | 0.758 |

**Supplemental Table 7.** Table showing the results of a grid search of the parameters for tSNE (learning rate and perplexity) that was applied to the Hydra data at the PCA step. The AJD shown is between the PCA embedding of the transformed data and a two-dimensional tSNE embedding generated with the selected set of parameters.

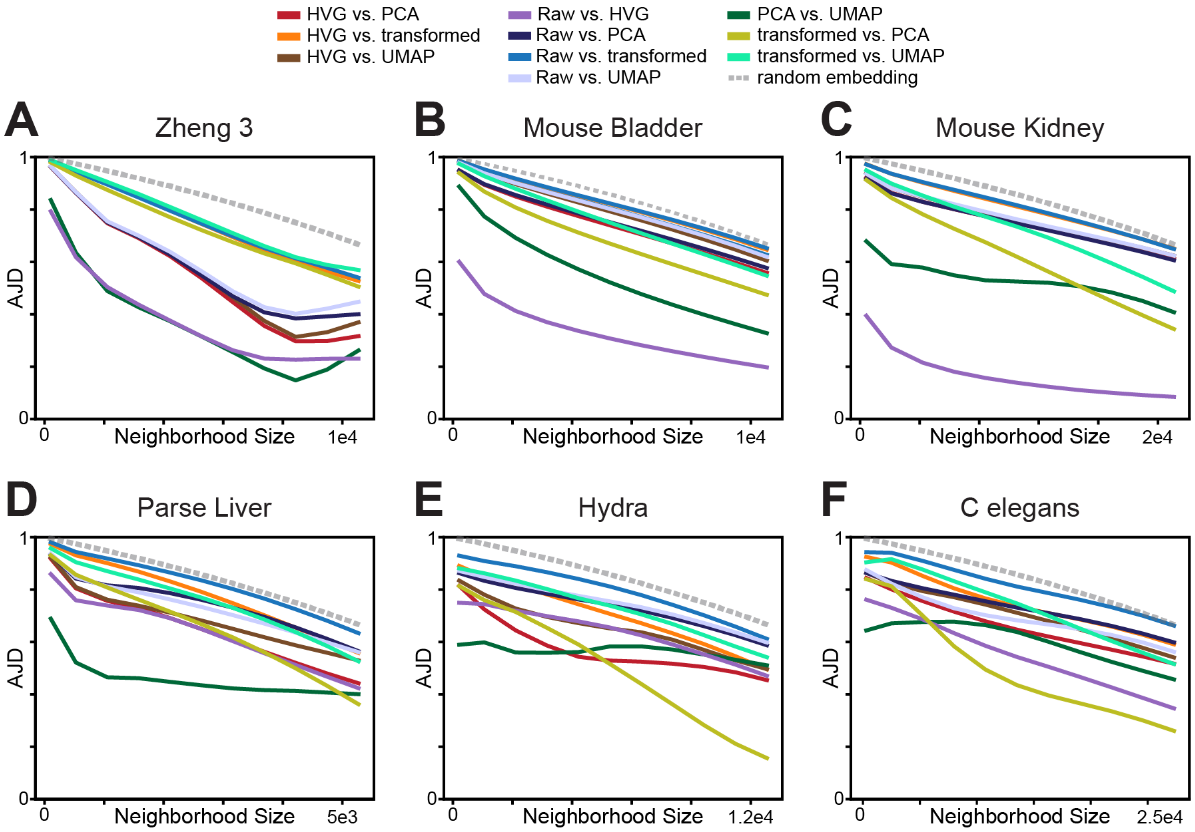


**Supplemental Figure 3.** Figure displaying the AJD as a function of neighborhood size control for each of the 10 Pairwise comparison between steps in the pipeline across all 6 datasets. These datasets are: **A.** Zheng 3 mixture of Monocytes, NK Cells and B-cells[29] **B.** Mouse Bladder cells[30] **C.** Mouse Kidney cells[31] **D.** Parse Bio Liver cells[32] **E.** Hydra cells[33] **F.** Developing *C. Elegans* cells[34] In addition to a pairwise comparison, a dotted line showing the Expected AJD given an embedding with randomized neighborhoods termed random overlap (See Supplementary Proof 2 for derivation) is displayed. Together they show that while each of the steps do preserve the data's topology better than randomly permuting the neighborhoods, the neighborhood size needed to reduce the data to have an AJD of less than 0.2 is often closer to fifty percent of the number of cells available in the dataset, a very weak definition of locality.

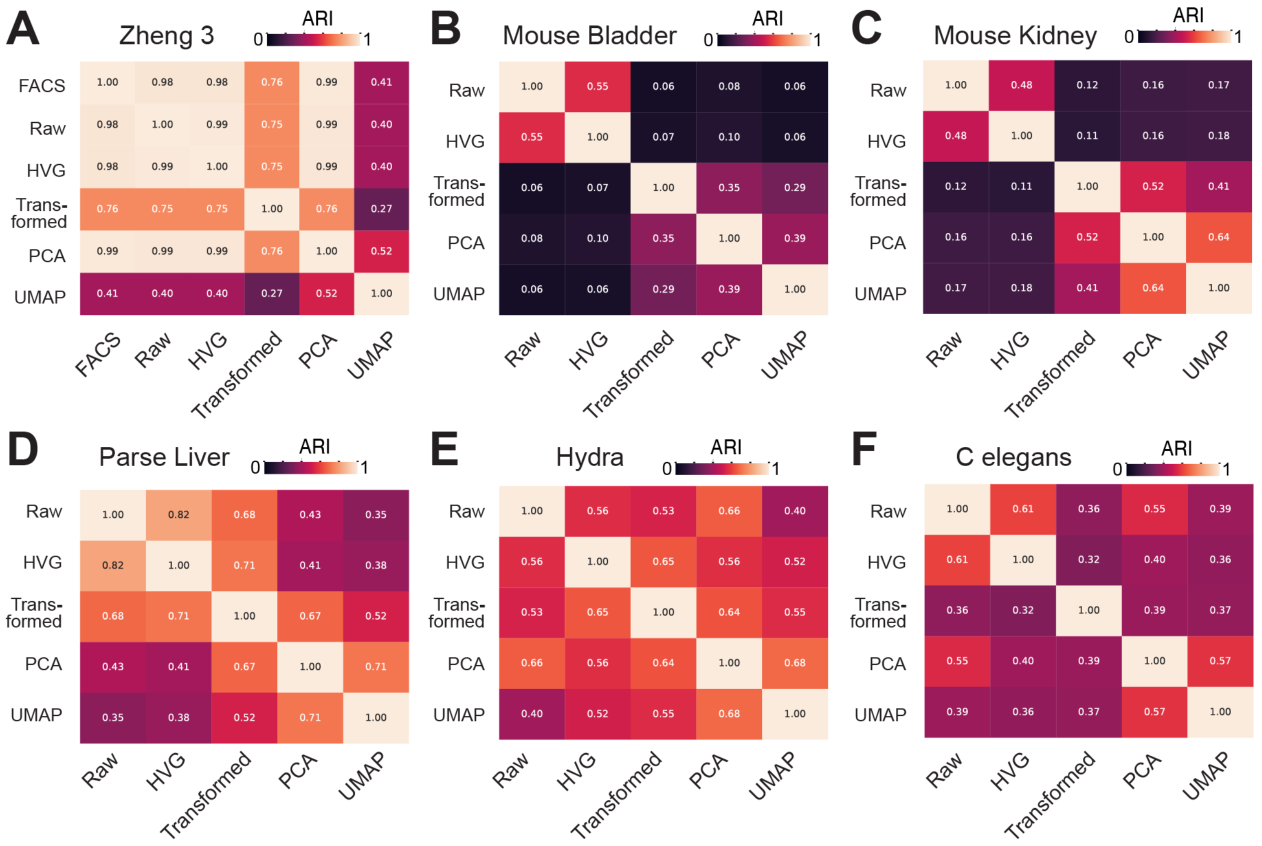


**Supplemental Figure 4.** Pairwise comparison of Louvain clustering for the six listed datasets analyzed. These datasets are **A.** Zheng 3 mixture of Monocytes, NK Cells and B-cells[29] **B.** Mouse Bladder cells[30] **C.** Mouse Kidney cells[31] **D.** Parse Bio Liver cells[32] **E.** Hydra cells[33] **F.** Developing *C. Elegans* cells[34] The ARI reported in each cell of the heatmap is the highest ARI reported between the Louvain clustering of the two different spaces as the resolution parameter was titrated.

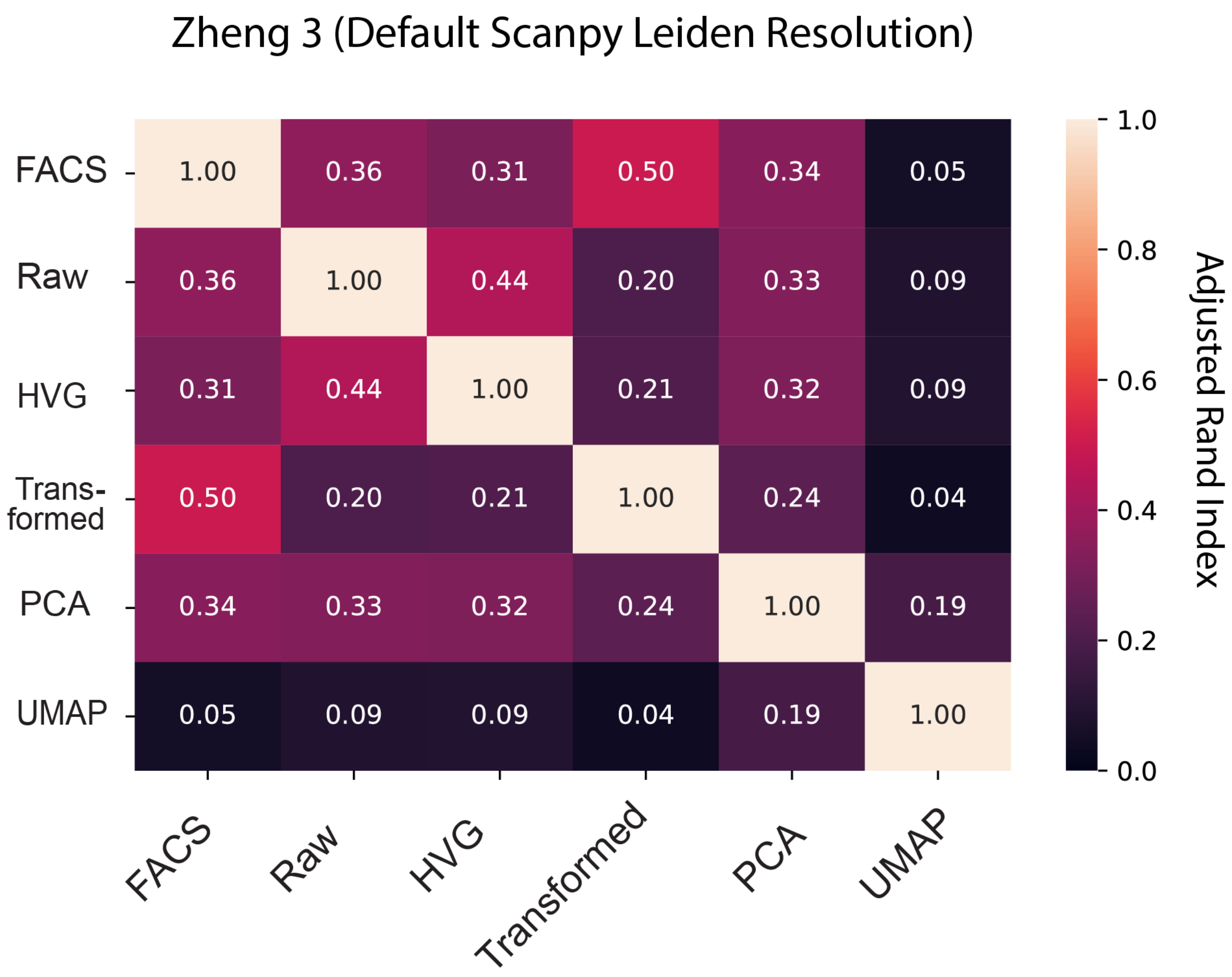


**Supplemental Figure 5.** Pairwise heatmap showing the ARI between the clusterings done on each of the different steps of the scRNA-seq pipeline for the Zheng 3 data. In this case, the Leiden clustering used the default resolution value, indicating that in a purely unsupervised sense, the distortion introduced in the pipeline and as measured by the AJD does cause massive changes in the local topology that can affect clustering results and adjusting the resolution to compensate can only mitigate this effect in certain datasets, particularly in ones where the composition of cells is well established
