## Supplemental Derivation for "A novel metric reveals previously unrecognized distortion in the analysis of scRNA-seq data"

### 1 Approximate AJD for random neighborhoods

Let $D$ be a dataset containing $N$ points in a $p$-dimensional dataset, i.e. $D \subset \mathbb{R}^p$. In the case of scRNA-seq data, this would correspond to $N$ cells with expression data for $p$ genes or features. Regardless, each point in dataset $D$ is just a vector $d \in \mathbb{R}^p$. Let there also be an embedding $E$ of that dataset containing $N$ cells and $q$ dimensions, with $q \leq p$. Note that, for every point in $d \in D$ there is a single corresponding point $e \in E$ that represents *the same* point in the lower-dimensional embedding. We can explicitly define this as a bijective function $f : D \to E$ that maps each point $d$ to the lower-dimensional representation of that point $f(d) \in E$.

For any point $d \in D$, define the set $A_d$ that contains the $k^{\text{th}}$ nearest neighbors of $d$. Similarly, for the corresponding point $f(d) \in E$, define the set of $k^{\text{th}}$ nearest neighbors in the low-dimensional representation as $B_d$; note here we use the subscript $d$ rather than $f(d)$ for notational convenience.

To calculate the AJD, we first calculate the Jaccard distance between these sets:

$$J_d = \frac{|A_d \,\Delta\, B_d|}{|A_d \cup B_d|} = 1 - \frac{|A_d \cap B_d|}{|A_d \cup B_d|},$$

where $|X|$ denotes the cardinality of the set $X$, $\Delta$, $\cup$ and $\cap$ are the symmetric difference, union and intersection operators on sets, respectively. To calculate the AJD, we average $J_d$ over all points $d \in D$.

We now have a simple model where we construct these lower-dimensional neighborhoods purely at random. To do this, we stipulate that $B_d$ is constructed by uniform sampling without replacement, with the constraint $|B_d| = |A_d| = k$.

Now define $l_d \equiv |A_d \cap B_d|$; by definition this means that $|A_d \cup B_d| = 2k - l_d$. Under the random sampling described above, it is clear that $l_d$ will be drawn from a hypergeometric distribution. In other words, we are constructing $B_d$ by sampling from $N$ total points without replacement, so $\Pr(l_d) \sim \text{Hypergeometric}(N, k, k)$. Here, we consider a "success" to be when one of the members of $A_d$, i.e. one of $d$'s original neighbors, is chosen to be $B_d$ through the random sampling process. There are thus a total of $k$ successes in the original population of $N$ points, and we perform $k$ trials because we are constructing $B_d$ so that it has $k$ elements.

The expectation of this value $l_d$ under this model is given by:

$$\mathrm{E}[l_d] = k\frac{k}{N} = \frac{k^2}{N}.$$

The Jaccard Distance between the sets $A_d$ and $B_d$ in this case is just:

$$J_d = 1 - \frac{l_d}{2k - l} = \frac{2k - 2l_d}{2k - l_d},$$

where we have just substituted $k$ and $l_d$ as appropriate.

We now make the approximation:

$$\text{AJD} = \mathrm{E}[J_d] = \mathrm{E}\left[\frac{2k - 2l_d}{2k - l_d}\right] \approx \frac{\mathrm{E}[2k - 2l_d]}{\mathrm{E}[2k - 2l_d]}.$$

Given this approximation we have:

$$\begin{aligned}
\mathrm{E}[J_d] &\approx \frac{2k - \mathrm{E}[l_d]}{2k - \mathrm{E}[l_d]} \\
\mathrm{E}[J_d] &\approx \frac{2k - 2\frac{k^2}{N}}{2k - \frac{k^2}{N}} \\
\mathrm{E}[J_d] &\approx \frac{2 - 2\frac{k}{N}}{2 - \frac{k}{N}} \\
\mathrm{E}[J_d] &\approx \frac{2N - 2k}{2N - k},
\end{aligned}$$

which is the approximation we use for this random model in the figures in the main text.
